# Localized, time-dependent responses of rat cranial bone to repeated mild traumatic brain injuries

**DOI:** 10.1101/2021.12.12.472300

**Authors:** Larissa K. Dill, Natalie A. Sims, Ali Shad, Chidozie Anyaegbu, Andrew Warnock, Yilin Mao, Melinda Fitzgerald, Bridgette D. Semple

## Abstract

While it is well-established that bone responds dynamically to mechanical loading, the effects of mild traumatic brain injury (mTBI) on cranial bone composition are unclear. We hypothesized that repeated mTBI (rmTBI) would change the microstructure of cranial bones, without gross skull fractures. To address this, young adult female Piebald Viral Glaxo rats received sham, 1x, 2x or 3x closed-head mTBIs delivered at 24h intervals, using a weight-drop device custom built for reproducible impact. Skull bones were collected at 2 or 10 weeks after the final injury/sham procedure, imaged by micro computed tomography and analyzed at predetermined regions of interest. In the interparietal bone, proximal to the injury site, modest increases in bone thickness was observed at 2 weeks, particularly following 3x mTBI. By 10 weeks, 2x mTBI induced a robust increase in the volume and thickness of the interparietal bone, alongside a corresponding decrease in the volume of marrow cavities in the diploë region. In contrast, neither parietal nor frontal skull samples were affected by rmTBI. Our findings demonstrate time- and location-dependent effects of rmTBI on cranial bone structure, highlighting a need to consider microstructural alterations to cranial bone when assessing the consequences of rmTBI.

## INTRODUCTION

It is increasingly recognized that repeated head impacts can have detrimental consequences on brain function. Mild traumatic brain injuries (mTBIs), including concussions, are the most common form of brain injury, and accumulating evidence suggests that repeated mTBIs (rmTBI) may result in persistent symptomology and chronic neuropathological effects, associated with a range of neuropsychiatric and neurodegenerative disorders ^1,2^. However, little consideration has been afforded to the direct or indirect effects of rmTBI on the cranial skull bones that protect the brain. Such knowledge is critical for understanding how the meninges, cerebral vasculature, lymphatics system and brain parenchyma also respond to a single injury, as well as any subsequent injuries ^3,4^.

Bone is a dynamic living tissue with a high capacity for remodeling in response to mechanical forces as well as environmental factors, such as hormonal influences or changes in gravity ^5^. Mechanical loading—induced by high-intensity exercise, for example, or trauma—is a well-established regulator of long bone mass, density and composition ^6–8^. However, how this phenomenon applies in the context of head impacts to skull bones has been poorly studied to date.

Cranial bone is a three-layered sandwich-like structure comprised of outer layers of compact cortical bone surrounding a central layer of irregular porous bone. This central diploë region is highly variable and responsive to applied mechanical forces ^9^. Significant damage to the cranium from a moderate or severe TBI, resulting in a skull fracture, has been associated with a greater degree of neuroinflammation and poorer functional outcomes in experimental TBI models ^10,11^. Similarly, the occurrence of a skull fracture in patients with severe TBI is a predictor of in-hospital mortality as well as unfavorable outcomes by 6 months post-injury ^12–14^ On the milder end of the injury spectrum, in the absence of skull fracture, mechanical loading may also influence bone structure, particularly in the context of repetitive impacts such as those sustained during competitive sporting activities.

Several recent studies have demonstrated that a TBI can promote bone formation, such as the development of neurogenic heterotopic ossification even in remote skeletal locations ^15–17^. Case studies have also reported that the skull periosteum, the membranous tissue that covers the bone surfaces, has considerable potential for osteogenesis in response to local hematomas ^18,19^. These findings suggest a complex relationship between bone and brain injury that likely extends beyond responsiveness to the direct mechanical forces applied by a head impact.

Preclinical studies have the potential to allow better understanding of the relationship between TBI and bone remodeling post-injury. In adolescent mice, we have previously reported that a single mTBI led to increased bone volume and strength of the impacted parietal bone by 5 weeks later, which may account for a reduced incidence of skull fractures when animals were exposed to a second mTBI impact in adulthood ^11^. Subsequently, another study found that mTBI in adult mice led to decreased bone porosity in the contralateral (uninjured) hemisphere, which appeared to be mediated by the cannabinoid-1 receptor ^20^. Together, this scant evidence drove the formation of our current aim: to evaluate whether rmTBI influences skull bone composition and structure.

A reproducible closed-head injury model was used to mimic single or rmTBI, as previously described ^21,22^. This model generates a mild brain injury in the absence of macroscopic brain damage or overt skull fractures. Previous work with this paradigm has demonstrated acute oxidative damage after rmTBI spaced 24 h apart, particularly after 2x mTBI, which worsens over weeks and months post-injury alongside microglial reactivity and indications of myelin pathology ^22,23^. Here, we turned our attention to the effects of head injury on the skull bones, hypothesizing that rmTBI would result in greater effects compared to a single mTBI, and those effects would develop in a time- and location-dependent manner.

## MATERIALS AND METHODS

### Animals and Ethics

Young adult female Piebald Viral Glaxo rats (12 weeks of age; 160-200 g) were sourced from the Animal Resource Center (Murdoch, WA, Australia). Animals were housed in standard cages under specific pathogen-free conditions and a 12:12 h light-dark cycle, with food and water available *ad libitum*, and acclimatized to housing conditions for at least one week prior to experiments. All procedures were approved by The University of Western Australia Animal Ethics Committee (#RA/3/100/1366) and conducted according to the Australian Code of Practice for the Care and Use of Animals for Scientific Purposes from the National Health and Medical Research Council (NHMRC) and in compliance with the ARRIVE guidelines.

### Experimental mTBI model

A reproducible closed-head injury model was used to mimic mild, repeated TBI, as previously described ^21,22^. In brief, rats were anesthetized with inhalant 4% isoflurane (4 L/min in oxygen) and maintained at 2% isoflurane (2 L/min oxygen), then positioned prone on a suspended Kimwipe (Kimberley-Clark, Irving, TX, USA). Using a custom-built weight drop device (Northeast Biomedical Inc., MA, USA) ^22–24^, injury was induced by release— down a guide tube—of a 250 g weight from 1 m height, to impact the head at the midline, 2-3 mm anterior to the front of the ears (i.e. aligned with Lambda on the underlying skull). Upon impact, the rat fell through the Kimwipe onto the foam mat below, allowing for rotational forces to be sustained, consistent with mechanisms common in human mTBI ^25–27^. Carprofen at 4 mg/kg i.p. (Norbrook Laboratories, Tullamarine, VIC, Australia) was administered postprocedure to all animals for analgesia. Rats were then allowed to recover on a heat pad at 37 °C until ambulatory.

Animals were randomized to one of four experimental groups, receiving either the sham procedure, 1x, 2x or 3x mTBI impacts, delivered at 24 h intervals. The 24 h inter-injury interval was chosen to approximate weekly exposure to sports-related rmTBI, based on evidence that biochemical processes such as basal metabolism occur around 6 times faster in rats than in humans ^28^. Importantly, rats that received 1 or 2 mTBIs underwent the sham procedure when mTBI was not administered, to ensure equivalent anesthesia exposure for all animals. The sham procedure was identical to that described above, except for the weight drop impact itself (Figure 1).

**Figure 1:**
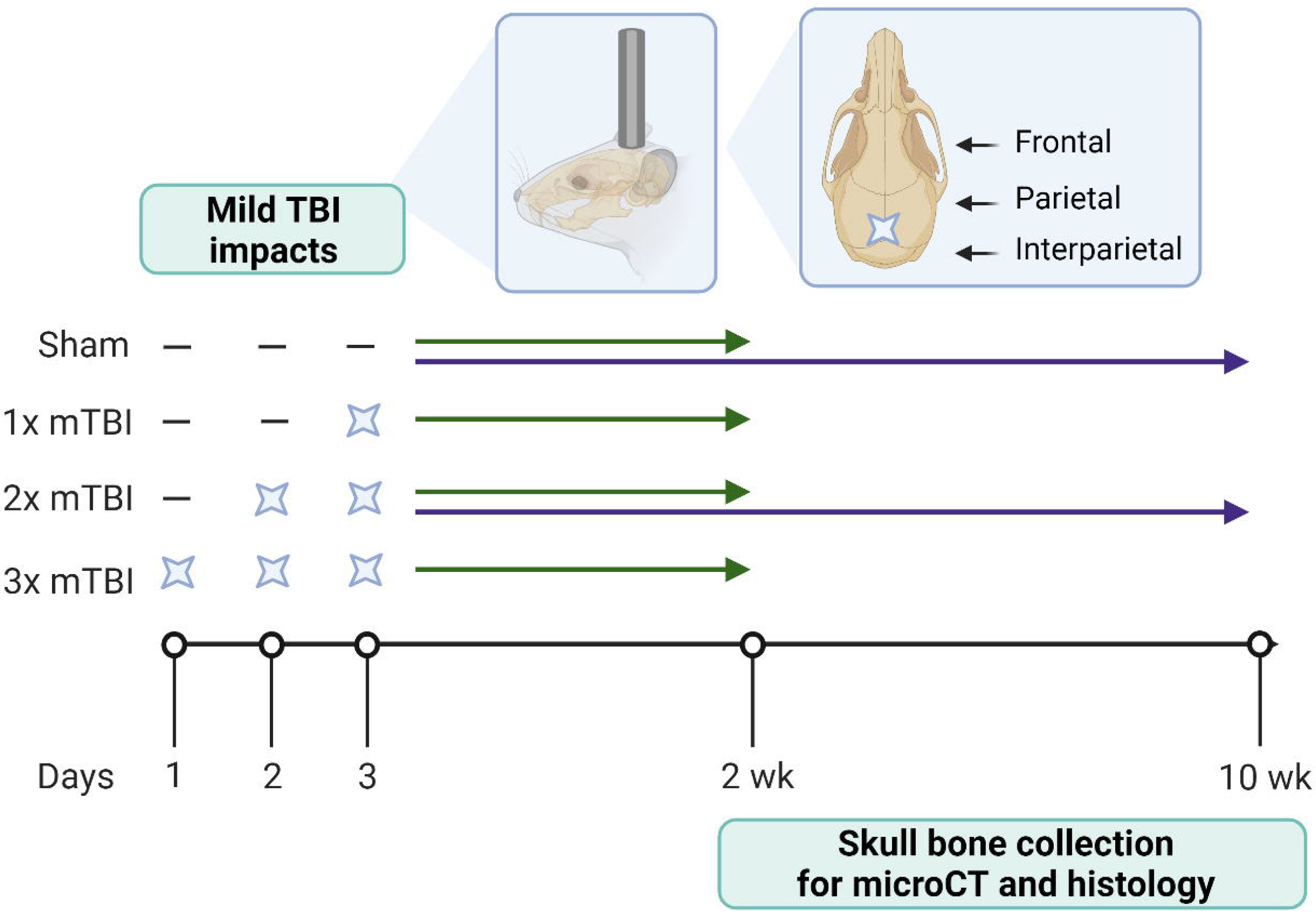
Schematic of study timeline. Rats received 0-3 x mTBI impacts or sham procedures spaced 24 h apart. Animals were euthanized at either 2 weeks (sham, 1x mTBI, 2x mTBI and 3x mTBI groups) or 10 weeks (sham and 2x mTBI groups), with the time point referring to time following the last mTBI/sham procedure for collection of skull bones for microCT and histology. Illustrations depict the site of impact of the weight drop model on the rat skull, from a sagittal and dorsal view, respectively. Time points were chosen based upon previous findings in this model, whereby 2x mTBI animals often showed the greatest effects compared to sham controls ^22,23^. Created with Biorender.com

### Sample collection

At 2 weeks following the final injury/sham procedure, a subset of animals were euthanized with 100 mg/kg i.p. sodium pentobarbital (Lethobarb, Virbac, Australia) and perfused transcardially with sterile saline (0.9% sodium chloride) followed by 4% paraformaldehyde in phosphate buffer (0.1 M, pH 7.2). Cranial bone samples were dissected and collected from rats from the following experimental groups: Sham, n=10; 1x mTBI, n=5; 2x mTBI, n=7; and 3x mTBI, n=5.

At 10 weeks after the final injury/sham procedure, cranial bones were collected from two groups: Sham (n=5) and 2x mTBI (n=5). The 2x mTBI group was chosen since previous findings from this model indicated that the 2x mTBI resulted in the greatest injury response, i.e. a higher degree of oxidative stress, microglial reactivity, and myelin pathology compared to a single mTBI or even the 3x mTBI group ^22,23^.

Collected samples were fixed in 4% paraformaldehyde then transferred into 70% ethanol for transport and storage.

### Micro computed tomography (microCT)

The rat cranial samples were imaged using a Siemens Inveon PET-SPECT-CT small animal scanner in micro-CT mode, at 80 kV and 500 uA with an exposure time of 600 ms (Monash Biomedical Imaging, Clayton, VIC Australia). Scans captured isotropic voxels of 9.7 μm in a field of approximately 1350 x 1350 x 1800 voxels.

Image stacks were reconstructed with x, y and z slice depth dimensions of 1 vx. ImageJ (https://imagej.nih.gov/ij/; National Institutes of Health, Bethesda, MD, USA) and the plugin BoneJ2^29^ were used on the high performance computing system MASSIVE^30^ to reconstruct, orient, crop and analyze the image stacks. All analyses were performed by an investigator blinded to experimental group.

Regions of interest (ROI) were defined relative to the midline and Lambda or Bregma, measured in voxels (Table 1). Interparietal and parietal bone ROI stacks consisted of 150 images, measuring 200 vx wide. Frontal bone ROI stacks consisted of 100 images, measuring 200 vx across. Bone and total tissue (including marrow space) was isolated using the statistical region merging segmentation ^31^ with the Otsu threshold algorithm ^32^ (thresholding settings described in *Supplementary Methods*). The BoneJ2 plugin was then used to measure bone volume in each sample, as well as the total tissue or object volume (i.e. bone plus marrow space) (see *Supplementary Methods*). The binary bone reconstructions were additionally used to calculate mean thickness of bone (averaged across 150 stack slices) and external surface area using BoneJ2.

**Table 1:**
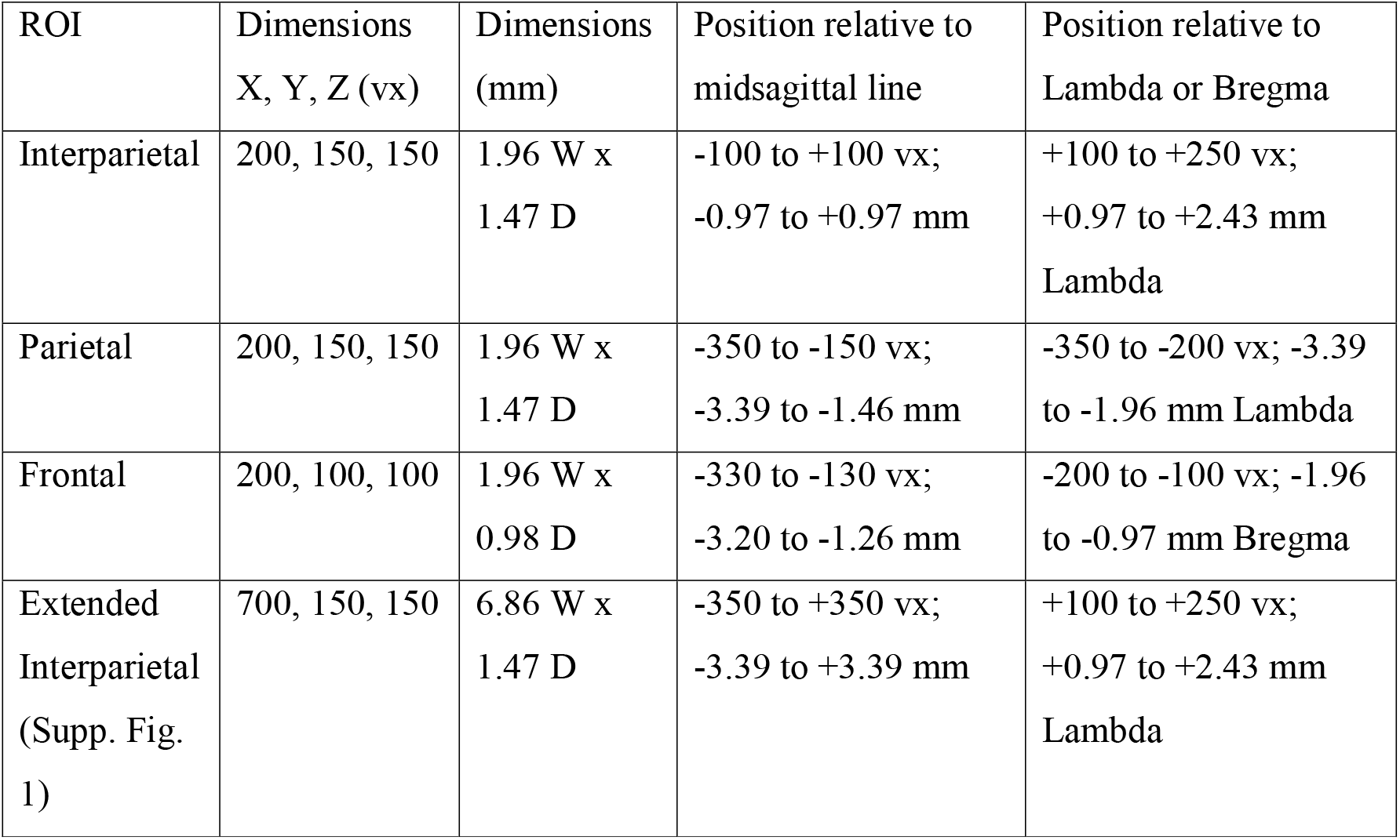
Region of interest (ROI) dimensions and loci.

Analysis of mean bone thickness along the rostrocaudal axis was performed using the BoneJ2 Slice Geometry tool. In order to validate the span of interparietal bone included in the primary analyses, the Slice Geometry tool was also used to interrogate mean thickness along the sagittal axis in extended interparietal ROIs consisting of 700 images, each measuring 150 vx deep.

The diploë component of interparietal and frontal bone ROIs was further investigated to quantify the marrow cavity volume of each sample. The diploë were isolated from interparietal and frontal cortical bone layers by limiting the sample height to 50 vx and 30 vx, respectively. The ImageJ plugin 3D Objects Counter ^33^ was then used to measure the frequency and volume of marrow cavities (i.e. void volume).

Both the left and right parietal bone from each bone sample was analyzed. As no differences were found in any measured parameters when comparing the left and right parietal bone regions, the left side only has been reported here for clarity. Frontal bone was only available for analysis in the 10 week samples, and the left side only was analyzed. One frontal bone sample was excluded due to the scan being of inappropriate resolution for analysis.

### Histology

Decalcification, sectioning and staining of bone samples was performed by the Monash Histology Platform (Alfred Research Alliance Node, Prahran, VIC, Australia). A 10% ethylenediaminetetraacetic acid (EDTA) solution (pH 7.4) was prepared and used to decalcify bone samples. A 2-3 week period was required to ensure complete decalcification, with EDTA solution being changed weekly. Samples were processed with Leica Peloris II tissue processor and then embedded in paraffin molds for microtomy using a Leica HistoCore BIOCUT microtome (Leica Microsystems, Wetzlar, Germany). ROIs for the left parietal and interparietal bone were defined relative to the Lambda suture. Five μm sections were collected at intervals of 150 μm in the range of −3.5 mm to −2.0 mm (parietal) and +1.0 mm to +2.5 mm (interparietal) for histological investigation. Standard hematoxylin and eosin (H&E) staining was performed on two sections per ROI using a Leica Autostainer XL. In brief, deparaffinized and rehydrated sections were incubated in Harris’s Hematoxylin for 5 min, rinsed and then dipped in acid alcohol solution. Following further rinsing, sections were incubated in eosin solution for 5 min, then processed for coverslipping.

### Statistical Analysis

MicroCT data were compiled and calculated using Microsoft Excel. Statistical tests were performed with GraphPad Prism software v. 9.0 (GraphPad Software Inc., La Jolla, CA). One-way analysis of variance (ANOVA) was used to evaluate differences between the four experimental groups at the 2 week time point. Tukey’s post-hoc tests were assessed when a significant main effect was detected by ANOVA. Unpaired t-tests were used to compare the sham and 2x mTBI group at the 10 week time point. Values are expressed as group mean ± SEM. Statistical significance was defined as *p* < 0.05, with *p* values and F statistics reported to 2 decimal places. For all graphical representation of data, both individual (grey dots) and mean group values (white and black bars) ± SEM are presented.

## RESULTS

### Proximal to the impact site, the interparietal bone shows increased thickness over time after repeated mTBI

The midline interparietal bone was firstly examined by microCT to assess the potential effects of a single or repeated mTBI to the young adult rat. At 2 weeks post-injury (Figure 2a-b), mTBI had a subtle but significant anabolic effect indicated by a greater bone volume (F_3,23_ = 4.27, *p* = 0.02) in the 3x mTBI group compared to the sham controls (Tukey’s post-hoc, *p* < 0.05). A slight increase in total object volume (i.e. comprising bone and marrow space) was also observed after TBI, but this did not reach statistical significance (Figure 2c; F_3,26_ = 2.55, *p* = 0.08); whereas external surface area was not different between groups (Figure 2d, F_3,23_ = 0.11, *p* = 0.42). Mean bone thickness (Figure 2e-f) was the main contributor to the increase in bone volume, with an increase after TBI, specifically in the 3x mTBI group (F_3, 23_ = 3.92, *p* = 0.02; post-hoc *p* < 0.05).

**Figure 2:**
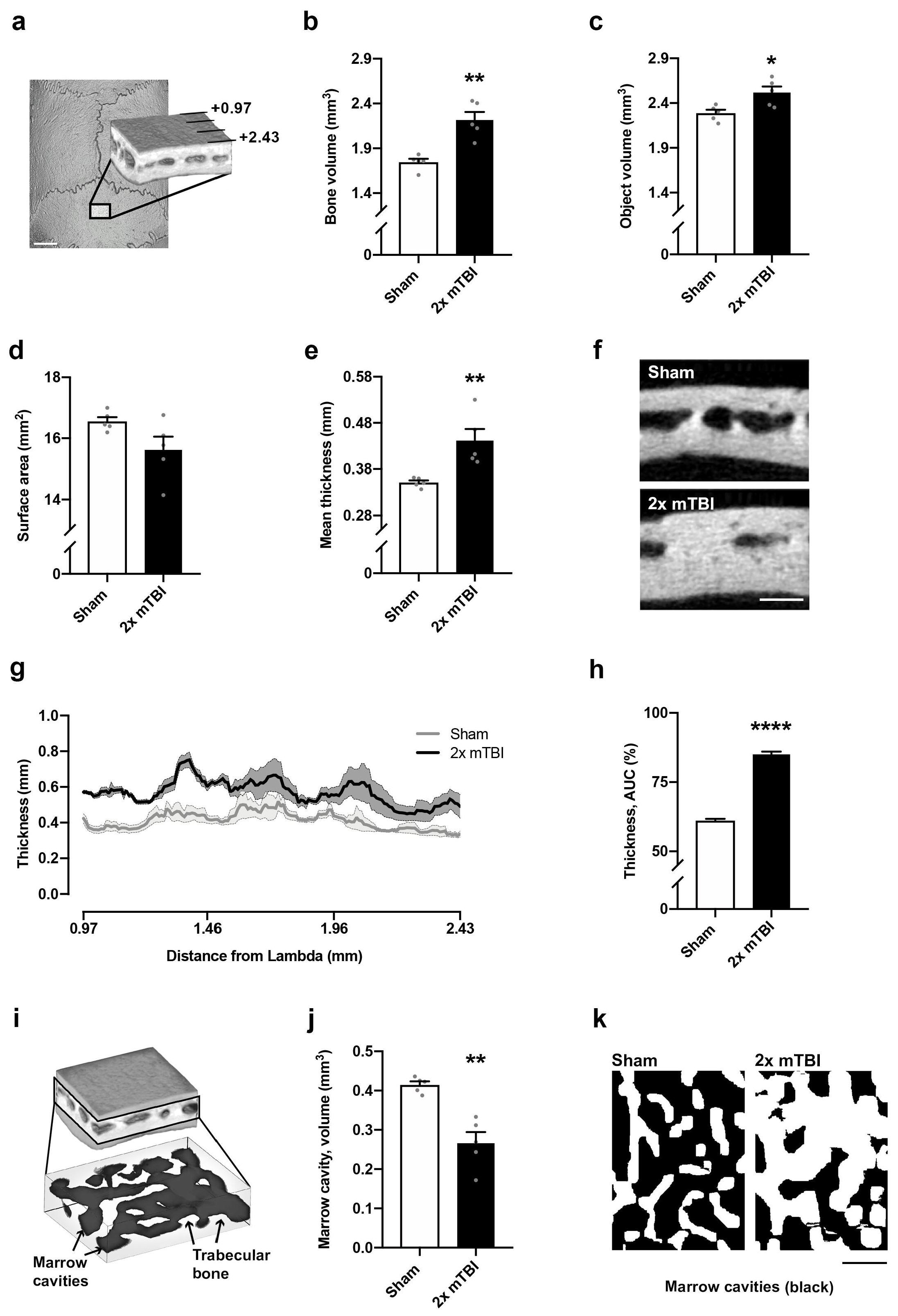
Modest increases in bone volume and thickness of the interparietal bone two weeks after rmTBI. The interparietal bone region-of-interest is depicted in (a), which consisted of 150 microCT images sampled at the midline, caudal to the Lambda suture margin (scale bar = 2 mm). Quantification of mineralized bone volume (b) and mineralized bone thickness (e) revealed an increase in 3x mTBI mice compared to sham controls (post-hoc, *p < 0.05); whereas no significant differences were observed between groups in total object volume (c) or surface area (d). Representative Sham and 3x mTBI greyscale microCT images (f) illustrate modest bone thickening with rmTBI (scale bar = 500 μm). In (g), bone marrow cavities (dark grey) were isolated from trabecular bone (light grey) in the diploë region of the interparietal bone scan ROIs for volumetric analysis (h), revealing a reduction in marrow cavity volume (one-way ANOVA p = 0.02; post-hoc n.s.). Representative superior-view, binary projections of the diploë (i) illustrate a subtle reduction in marrow cavities with injury (scale bar = 500 μm; black indicates marrow cavity and white indicates bone). N = 5-10/group; one-way ANOVAs with Tukey’s post-hoc as appropriate.

Repeated mTBI also appeared to reduce the total volume of bone marrow cavities in the diploë region at 2 weeks post-injury (Figure 2g-i), with a significant effect of injury detected by one-way ANOVA (F_3,23_ = 4.03, *p* = 0.02), but post-hoc analysis did not reveal a significant reduction in the 2x mTBI and 3x mTBI groups compared to sham controls (post-hoc; *p* = 0.09 and *p* = 0.06, respectively).

At 10 weeks post-injury, these anabolic effects on the interparietal bone were more pronounced. Specifically, we observed a greater bone volume (Figure 3a-b; t_8_ = 4.88, *p* < 0.01) and total object volume (bone plus marrow cavity space; Figure 3c; t_8_ = 3.00, *p* = 0.02) in 2x mTBI mice compared to sham controls, but no difference in the surface area between groups (Figure 3d; t_8_ = 2.04, *p* = 0.08).

**Figure 3:**
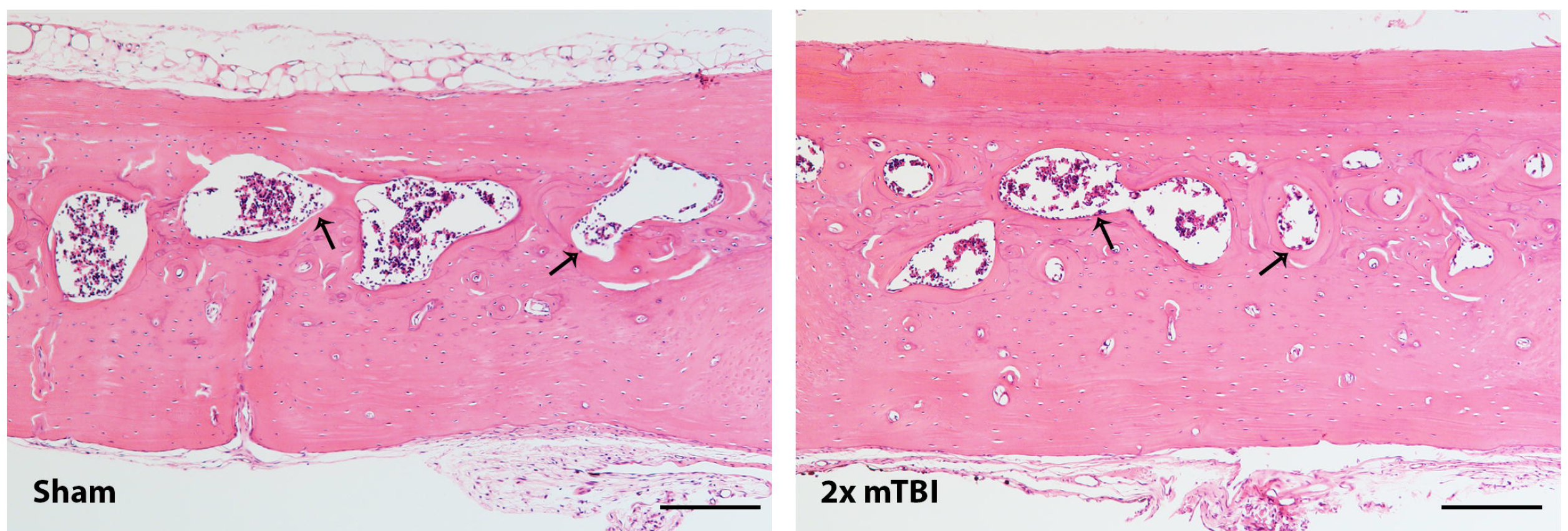
A robust increase in volume and thickness of the interparietal bone was observed at 10 weeks after rmTBI. The interparietal bone region-of-interest (ROI) is depicted in (a), mirroring the ROI considered at the 2 week time point (scale bar = 2 mm). Quantification of mineralized bone volume (b) revealed a significant increase in the 2x mTBI group compared to sham controls (t-test, **p < 0.01), as well as the object volume (i.e. bone plus marrow cavity space; c; *p = 0.02). Exterior surface area did not reach statistical significance (d). Mean mineralized bone thickness, averaged across 150 adjacent sections, was significantly increased in 2x mTBI mice (e, post-hoc **p < 0.001), as visualized by representative greyscale microCT images from the ROI z-stack center (f, scale bar = 500 μm). Mineralized bone thickness was also visualized across the rostrocaudal axis (g), with the 2x mTBI group (black line, dark grey error range) across the 150-image span of the 1.47 mm deep interparietal ROI, which was quantified by area under curve analyses (h, ****p < 0.0001). The volume of marrow cavities (dark grey) were distinguished from trabecular bone (light grey) in the diploë region of the interparietal bone (i), revealing a significant decrease in total cavity volume in the 2x mTBI group compared to sham controls (j, **p < 0.01). Representative superior view, binary projections of the diploë (k) illustrates a robust reduction in marrow cavities in 2x mTBI mice compared with sham controls (scale bar = 500 μm). N = 5/group; unpaired student’s t-tests.

Mean bone thickness at 10 weeks post-injury, when averaged across all 150 adjacent microCT sections (Figure 3e; t_8_ = 3.50, *p* = 0.01), was significantly greater in 2x mTBI mice compared to sham (** *p* < 0.001). To evaluate whether this effect varied across the rostrocaudal axis, bone thickness was also depicted across the ROI cross-section, which was quantified by area under the curve analysis to confirm a significant increase in bone thickness in the 2x mTBI group compared to sham (Figure 3g-h; t_1600_ = 21.75, *p* < 0.0001). Evaluation of interparietal bone thickness across a sagittal cross-section demonstrated that this effect was most prominent close to the midline and site of TBI impact (<u>Suppl. Figure 1</u>).

Also at 10 weeks post-injury, the volume of bone marrow cavities was significantly reduced in 2x mTBI mice compared to sham controls (Figure 3i-k; t_8_ = 5.02, *p* < 0.01), with this quantified difference being clearly evident upon visual inspection of representative images. Together, these findings indicate greater interparietal diploë trabecular (strut) bone volume and thickness, and a corresponding reduction in marrow cavities that progresses from 2 to 10 weeks post-injury in mice that sustained 2x mTBI insults.

These TBI-induced changes in skull bone thickness and the reduction in marrow cavities within the diploë region were also apparent by H&E staining of decalcified bone sections (Figure 4), whereas no overt differences were observed between H&E-stained bone of sham and rmTBI animals when collected at the 2 week time point (not shown).

**Figure 4:**
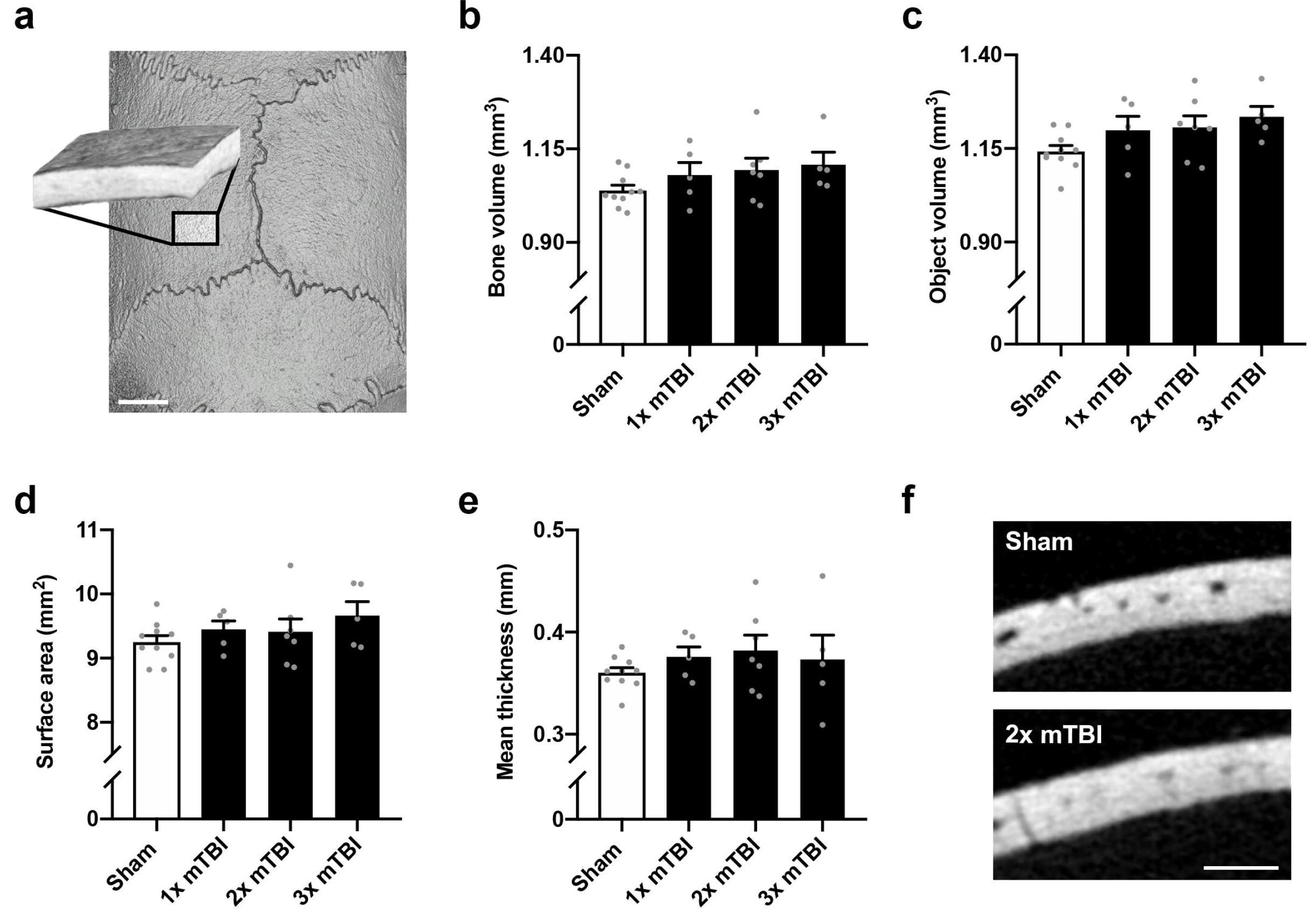
Hematoxylin and eosin staining of representative interparietal skull bone samples from sham and 2x mTBI rats, at 10 weeks post-injury. A larger volume of trabecular bone is evident in the 2x mTBI sample compared to sham, alongside a reduction in the volume of marrow cavities (arrows). Scale bar = 200 μm.

### Distal to the impact site, the parietal and frontal bones were unaffected by repeated mTBI

We next examined the mid-parietal bone by microCT (Figure 5a), as a region more distal from the injury impact site. At 2 weeks post-injury, there was no significant effect of injury on bone volume (Figure 5b; F_3,23_ = 1.53, *p* = 0.23), total object volume (Figure 5c; F_3,23_ = 2.45, *p* = 0.09) or exterior surface area (Figure 5d; F_3,23_ = 1.14, *p* = 0.35). In addition, bone thickness was not affected by injury at this site (Figure 5e-f; F_3,32_ = 0.67, *p* = 0.58).

**Figure 5:**
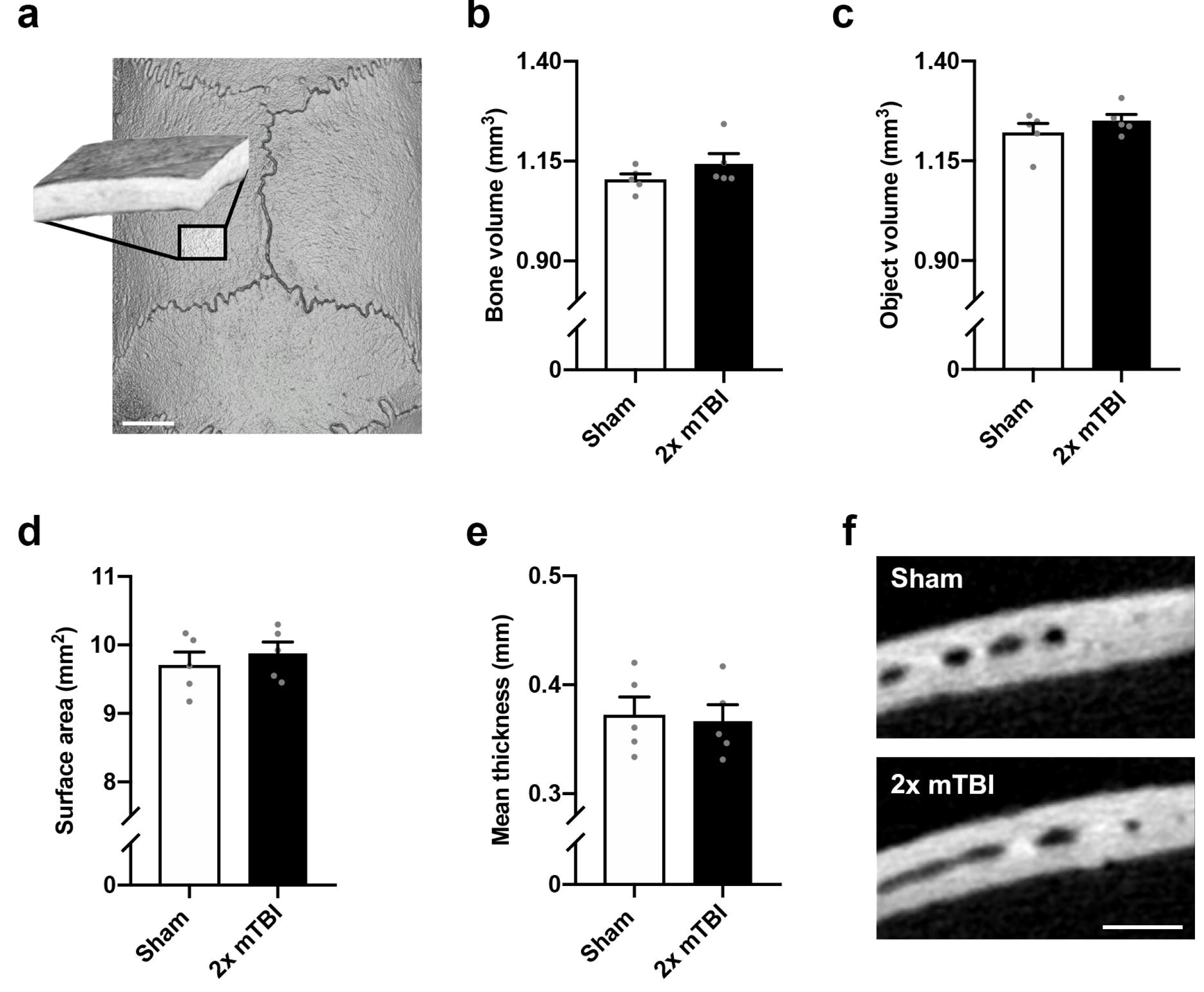
No change in volume, surface area or thickness of the parietal bone at 2 weeks after rmTBI. The parietal bone region-of-interest (ROI) consisting of 150 microCT images (a) was sampled from the mediocaudal left parietal bone, adjacent to the midline and Lambda suture margins (scale bar = 2 mm). Neither single nor repeated mTBI influenced bone volume (b, p = 0.23), total object volume (c, p = 0.09), or surface area (d, p = 0.35). Mean bone thickness was also not affected by injury (e, p = 0.58), as demonstrated by representative greyscale microCT images from the ROI z-stack center (f, scale bar = 500 μm). Parietal ROIs from both hemispheres were examined but showed similar findings, hence only the left ROI is presented. N=5-10/group; one-way ANOVA.

At 10 weeks post-injury, the left parietal bone was similarly unaffected by repeated mild TBI, with no differences between sham and 2x mTBI groups in mineralized bone volume (Figure 6a-b; t_8_ = 1.33, *p* = 0.22), total object volume (<u>Figure 6c</u>; F_8_ = 1.05, *p* = 0.32), or exterior surface area (Figure 6d; F_8_ = 0.68, *p* = 0.52). Further, bone thickness was also not affected by injury (Figure 6e-f; F_8_ = 0.27, *p* = 0.80).

**Figure 6:**
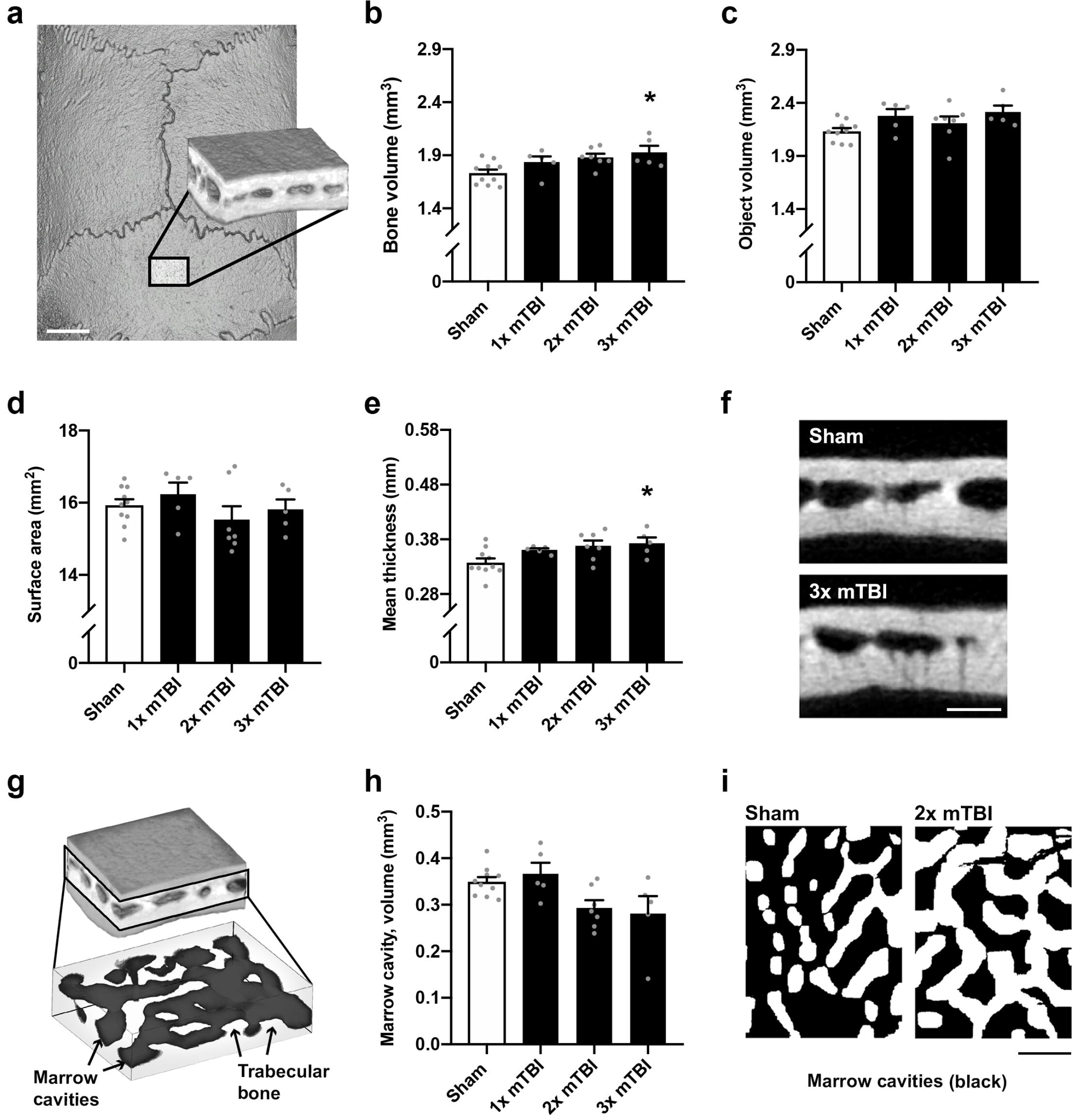
No change in volume, surface area or thickness of the parietal bone at 10 weeks after rmTBI. The parietal bone region-of-interest (ROI) consisting of 150 microCT images (a) was sampled from the mediocaudal left parietal bone, adjacent to the midline and Lambda suture margins (scale bar = 2 mm). Neither single nor repeated mTBI influenced mineralized bone volume (b, p = 0.22), total object volume (c, p = 0.32), or exterior surface area (d, p = 0.52). Mean mineralized bone thickness was also not affected by injury (e, p = 0.80), as demonstrated by representative greyscale microCT images from the ROI z-stack center (f, scale bar = 500 μm). Parietal ROIs from both hemispheres were examined but showed comparable findings; hence only the left ROI is presented. N=5-10/group; one-way ANOVA.

Finally, no differences in any quantified parameters (mineralized bone volume, object volume (bone plus marrow cavity space), exterior surface area, bone thickness, or marrow cavity volume) were observed in the left frontal bone at 10 weeks post-injury, even more remote to the impact site, in 2x mTBI samples compared to Sham controls (<u>Suppl. Figure 2</u>).

## DISCUSSION

In this study, we examined whether mild closed-head injuries to the female rat skull—up to three impacts, spaced 24 h apart—would result in changes within cranial bone detectable by microCT and histology. In this model of mTBI, we have previously reported that 2x mTBI animals showed modest cognitive memory deficits, increased microglial reactivity, lipid peroxidation and reduced white matter myelin integrity compared to sham controls and a single mTBI insult; although intriguingly, increasing the number of impacts beyond two did not necessarily exacerbate these effects ^22,23^. Now, we have extended our characterization of this model to reveal that rmTBI also modifies the impacted skull bone in a time- and location-dependent manner. Specifically, a localized increase in cranial bone thickness and volume, coincidental with a reduction in marrow cavity volume, appeared to evolve from 2 weeks to 10 weeks, and was restricted to the interparietal bone in close proximity to the impact location. These findings confirm our previous pilot work suggesting that a single mTBI over the unilateral parietal bone in adolescent male mice led to a localized increase in bone thickness after several weeks ^11^, and indicate that this phenomenon can be observed across different species, experimental TBI models, and injury locations.

The most likely mechanism at play here is mechanical force, in alignment with our observation of altered bone structure in close proximity to the impact site, but not in more remote regions of the cranium. While mTBI or concussive head impacts in isolation typically involve mechanical forces below the threshold required to induce a skull fracture, mechanical loading to the cranium may nonetheless result in subtle and transient deformation of the skull, stimulating a localized reparative response characterized by enhanced bone formation by osteoblasts ^34^. Indeed, a large body of work has demonstrated this phenomenon in the context of tibial compression, where mechanical loading stimulates increased cortical and trabecular bone formation ^35^. Future studies incorporating sophisticated techniques to label and track the response and activity of osteoblasts and osteoclasts will be required to define the precise mechanisms involved. It is plausible, for example, that changes in osteocyte production of the mechanosensitive Wnt inhibitor sclerostin is involved in this response ^36–38^.

Although there is a scarcity of published literature on skull responses after mild head impacts, cranial bone thickening has previously been reported in the context of hydrocephalus, raising the intriguing possibility that abnormal pressure from cerebrospinal fluid build-up or swelling—as can occur after a TBI—may also act as an intracranial mechanical regulator of bone growth ^39^. Alternatively, as most of these hydrocephalus patients had ventricular shunts placed during early childhood resulting in chronic intracranial hypotension, the subsequent lack of outward pressure on the developing skull may result in bone thickening to fill the vacant space ^40–42^. It is unclear whether similar interactions might occur between the cranium and cerebrospinal fluid in the context of TBI, particularly after mTBI in which significant swelling or changes in intracranial pressure are unlikely. However, these case studies do highlight the potential of skull bone to adapt and respond to external stimuli.

The implications of our findings—that rmTBI resulted in localized thickening of the skull by 10 weeks post-injury—may prove controversial. Cranial bone thickness and strength is developmentally-dependent ^43^, and skull thickness is an important determinant of the propensity of the skull to deform and/or fracture ^44,45^. Even in adulthood, experimental closed head impacts to middle-aged (20 week old) mice result in a lower incidence of skull fracture compared to similar injuries in younger mice (10 weeks old), coinciding with the older animals having thicker skulls ^10^. Across a longer time course, cortical thinning, a key component of fracture risk, is associated with aging in most of the skeletal system, including the skull ^46^. Our findings raise the interesting question of whether increased skull thickness after repeated mTBI has consequences for the strength of the bone, and how a subsequent force is transduced. The mechanical strength of cranial bones were not evaluated in this study; such an investigation would require the development of validated methods, and would be a useful addition for future studies to determine the functional consequences of observed changes in bone thickness.

Theoretically, thicker cranial bone would translate to any subsequent head impacts being transduced to a lesser extent than in a skull with no prior impacts. Indeed, we observed this previously in the mouse; a prior mTBI at postnatal day 35 reduced the incidence of skull fracture after a second mTBI induced at postnatal day 70 ^11^. One might surmise from this logic that mild impacts to the skull, below the threshold required to induce a skull fracture, might be protective, in that they promote bone growth to provide greater protection against future injuries—a potentially useful phenomenon in contact sports and military environments in which there is a risk of repeated head impacts. However, we are certainly not advocating for voluntarily acquiring repeated head impacts as a form of preconditioning! Accumulating evidence continues to demonstrate that repeated exposure to head impacts is detrimental to brain structure and function, as well as recovery after subsequent hits ^47–50^. Rather, our findings highlight the need for studies to consider how the whole head is impacted by a mTBI or concussion, including the skull, as well as connective tissue, meninges, skin and vasculature; and to better understand the complex brain-bone relationship under conditions of repetitive mild mechanical loading.

## CONCLUSION

We herein report greater localized cranial thickness and lowered calvarial porosity after rmTBI in the female rat, which are both time- and location-dependent. While the consequences of such changes on the mechanical properties of the skull remain to be elucidated, our findings support future studies adopting a more holistic investigation of the mechanisms and consequences of repeated head impacts on the skull as well as the brain tissue itself.

## Data Availability Statement

The datasets generated during and/or analyzed during the current study are available from the corresponding author on reasonable request.

## Acknowledgements

This study was indirectly supported by funding to BDS in the form of a Career Development Fellowship (APP1141347) from the National Health and Medical Research Council of Australia (NHMRC), and an Establishment Grant from the Central Clinical School, Monash University. NAS is supported by an NHMRC Senior Research Fellowship (APP1154819). This study was supported by the MASSIVE high-performance computing facility (www.massive.org.au), and assistance from the Monash Histology Platform (Alfred Research Alliance and Clayton nodes). The authors also thank Dr. Rhys Brady and Dr. Stuart McDonald for early discussions regarding the project, and Dr. Michael De Veer and Tara Sepehrizadeh from the Monash Biomedical Imaging Facility for their technical assistance.

## Author Contributions

Conceptualization: LKD, MF, BDS. Experimental investigation: LKD, CA, AW, YM. Methodologies, analysis and resources: LKD, NAS, AS, MF, BDS. Supervision, project administration, funding acquisition: MF, BDS. Manuscript writing: LKD, BDS. Manuscript editing: all authors. All authors have read and approved the manuscript for publication.

## REFERENCES

1. Huber, B.R., Alosco, M.L., Stein, T.D. & McKee, A.C. Potential Long-Term Consequences of Concussive and Subconcussive Injury. Physical medicine and rehabilitation clinics of North America 27, 503–511 (2016).

2. Hunter, L.E., Branch, C.A. & Lipton, M.L. The neurobiological effects of repetitive head impacts in collision sports. Neurobiol Dis 123, 122–126 (2019).

3. Stokely, M.E. & Orr, E.L. Acute effects of calvarial damage on dural mast cells, pial vascular permeability, and cerebral cortical histamine levels in rats and mice. J Neurotrauma 25, 52–61 (2008).

4. Bolte, A.C., et al. Meningeal lymphatic dysfunction exacerbates traumatic brain injury pathogenesis. Nature communications 11, 4524 (2020).

5. Crockett, J.C., Rogers, M.J., Coxon, F.P., Hocking, L.J. & Helfrich, M.H. Bone remodelling at a glance. J Cell Sci 124, 911–918 (2011).

6. Guglielmini, C., et al. Relationship between physical activity level and bone mineral density in two groups of female athletes. The quarterly journal of nuclear medicine: official publication of the Italian Association of Nuclear Medicine (AIMN) [and] the International Association of Radiopharmacology (IAR) 39, 280–284 (1995).

7. Buckwalter, J.A. & Cooper, R.R. Bone structure and function. Instructional course lectures 36, 27–48 (1987).

8. Bergmann, P., et al. Loading and skeletal development and maintenance. J Osteoporos 2011, 786752–786752 (2010).

9. McElhaney, J.H., et al. Mechanical properties on cranial bone. Journal of biomechanics 3, 495–511 (1970).

10. Zvejniece, L., et al. Skull Fractures Induce Neuroinflammation and Worsen Outcomes after Closed Head Injury in Mice. J Neurotrauma 37, 295–304 (2020).

11. McColl, T.J., et al. Mild Traumatic Brain Injury in Adolescent Mice Alters Skull Bone Properties to Influence a Subsequent Brain Impact at Adulthood: A Pilot Study. Front Neurol 9, 372 (2018).

12. Fujiwara, G., et al. Association of skull fracture with in-hospital mortality in severe traumatic brain injury patients. The American journal of emergency medicine 46, 78–83 (2021).

13. Tseng, W.C., et al. The association between skull bone fractures and outcomes in patients with severe traumatic brain injury. J Trauma 71, 1611–1614; discussion 1614 (2011).

14. Fabbri, A., Servadei, F., Marchesini, G., Stein, S.C. & Vandelli, A. Early predictors of unfavourable outcome in subjects with moderate head injury in the emergency department. J Neurol Neurosurg Psychiatry 79, 567–573 (2008).

15. Alexander, K.A., Tseng, H.W., Salga, M., Genêt, F. & Levesque, J.P. When the Nervous System Turns Skeletal Muscles into Bones: How to Solve the Conundrum of Neurogenic Heterotopic Ossification. Current osteoporosis reports 18, 666–676 (2020).

16. Anthonissen, J., Steffen, C.T., Hofmann, A. & Victor, J. The pathogenesis of heterotopic ossification after traumatic brain injury. A review of current literature. Acta orthopaedica Belgica 86, 369–377 (2020).

17. Brady, R.D., et al. Closed head experimental traumatic brain injury increases size and bone volume of callus in mice with concomitant tibial fracture. Scientific reports 6, 34491 (2016).

18. Choi, J., et al. Chronic ossified subperiosteal hematoma of the skull in an 11-year-old child: a case report. Child’s nervous system: ChNS: official journal of the International Society for Pediatric Neurosurgery 27, 1165–1168 (2011).

19. Wang, Y. & Zhang, J. Ossification of subperiosteal hematoma: the potential of periosteal osteogenesis in cranioplasty. The Journal of craniofacial surgery 24, 1603–1605 (2013).

20. Eger, M., et al. Bone Anabolic Response in the Calvaria Following Mild Traumatic Brain Injury is Mediated by the Cannabinoid-1 Receptor. Scientific reports 9, 16196 (2019).

21. Anyaegbu, C.C., et al. Simultaneous flow cytometric characterization of multiple cell types and metabolic states in the rat brain after repeated mild traumatic brain injury. J Neurosci Methods 359, 109223 (2021).

22. Yates, N.J., et al. Repeated mild traumatic brain injury in female rats increases lipid peroxidation in neurons. Exp Brain Res 235, 2133–2149 (2017).

23. Fehily, B., et al. Differential responses to increasing numbers of mild traumatic brain injury in a rodent closed-head injury model. J Neurochem 149, 660–678 (2019).

24. Mao, Y., et al. The Effects of a Combination of Ion Channel Inhibitors in Female Rats Following Repeated Mild Traumatic Brain Injury. Int J Mol Sci 19(2018).

25. Meehan, W.P.r., Zhang, J., Mannix, R., Whalen, M.J. Increasing recovery time between injuries improves cognitive outcome after repetitive mild concussive brain injuries in mice. Neurosurgery 71, 885–891 (2012).

26. Mannix, R., et al. Clinical correlates in an experimental model of repetitive mild brain injury. Ann Neurol 74, 65–75 (2013).

27. Mychasiuk, R., Hehar, H., Candy, S., Ma, I. & Esser, M.J. The direction of the acceleration and rotational forces associated with mild traumatic brain injury in rodents effect behavioural and molecular outcomes. J Neurosci Methods 257, 168–178 (2016).

28. Agoston, D.V. How to Translate Time? The Temporal Aspect of Human and Rodent Biology. Front Neurol 8, 92 (2017).

29. Domander, R., Felder, A.A. & Doube, M. BoneJ2 - refactoring established research software Wellcome Open Research 6, 37 (2021).

30. Goscinski, W.J., et al. The multi-modal Australian ScienceS Imaging and Visualization Environment (MASSIVE) high performance computing infrastructure: applications in neuroscience and neuroinformatics research. Frontiers in Neuroinformatics 8(2014).

31. Nock, R. & Nielsen, F. Statistical region merging. IEEE Transactions on Pattern Analysis and Machine Intelligence 26, 1452–1458 (2004).

32. Otsu, N. A Threshold Selection Method from Gray-Level Histograms. IEEE Transactions on Systems, Man, and Cybernetics 9, 62–66 (1979).

33. Bolte, S. & Cordelieres, F.P. A guided tour into subcellular colocalization analysis in light microscopy. J Microscopy 224, 213–232 (2006).

34. Castillo, A.B. & Leucht, P. Bone Homeostasis and Repair: Forced Into Shape. Current rheumatology reports 17, 58 (2015).

35. De Souza, R.L., et al. Non-invasive axial loading of mouse tibiae increases cortical bone formation and modifies trabecular organization: a new model to study cortical and cancellous compartments in a single loaded element. Bone 37, 810–818 (2005).

36. Robling, A.G., et al. Mechanical stimulation of bone in vivo reduces osteocyte expression of Sost/sclerostin. The Journal of biological chemistry 283, 5866–5875 (2008).

37. Tu, X., et al. Sost downregulation and local Wnt signaling are required for the osteogenic response to mechanical loading. Bone 50, 209–217 (2012).

38. Kang, K.S., et al. Loss of mechanosensitive sclerostin may accelerate cranial bone growth and regeneration. J Neurosurg 129, 1085–1091 (2018).

39. Huggare, J. & Rönning. Growth of the cranial vault: influence of intracranial and extracranial pressures. Acta odontologica Scandinavica 53, 192–195 (1995).

40. Lucey, B.P., March, G.P., Jr. & Hutchins, G.M. Marked calvarial thickening and dural changes following chronic ventricular shunting for shaken baby syndrome. Archives of pathology & laboratory medicine 127, 94–97 (2003).

41. Di Preta, J.A., Powers, J.M. & Hicks, D.G. Hyperostosis cranii ex vacuo: a rare complication of shunting for hydrocephalus. Human pathology 25, 545–547 (1994).

42. Villani, R., et al. Skull changes and intellectual status in hydrocephalic children following CSF shunting. Developmental medicine and child neurology. Supplement, 78–81 (1976).

43. Lillie, E.M., Urban, J.E., Lynch, S.K., Weaver, A.A. & Stitzel, J.D. Evaluation of Skull Cortical Thickness Changes With Age and Sex From Computed Tomography Scans. Journal of bone and mineral research: the official journal of the American Society for Bone and Mineral Research 31, 299–307 (2015).

44. Hubbard, R.P., Melvin, J.W. & Barodawala, I.T. Flexure of cranial sutures. Journal of biomechanics 4, 491–496 (1971).

45. Peterson, J. & Dechow, P.C. Material properties of the inner and outer cortical tables of the human parietal bone. The Anatomical record 268, 7–15 (2002).

46. Torimitsu, S., et al. Statistical analysis of biomechanical properties of the adult skull and age-related structural changes by sex in a Japanese forensic sample. Forensic Sci Int 234, 185.e181–189 (2014).

47. Pham, L., et al. Behavioral, axonal, and proteomic alterations following repeated mild traumatic brain injury: Novel insights using a clinically relevant rat model. Neurobiol Dis 148, 105151 (2021).

48. Gold, E.M., et al. Repeated Mild Closed Head Injuries Induce Long-Term White Matter Pathology and Neuronal Loss That Are Correlated With Behavioral Deficits. ASN neuro 10, 1759091418781921 (2018).

49. Bolton Hall, A.N., Joseph, B., Brelsfoard, J.M. & Saatman, K.E. Repeated Closed Head Injury in Mice Results in Sustained Motor and Memory Deficits and Chronic Cellular Changes. PLoS One 11, e0159442 (2016).

50. Mannix, R., et al. Chronic gliosis and behavioral deficits in mice following repetitive mild traumatic brain injury. J Neurosurg 121, 1342–1350 (2014).

