## Supplementary files for "Localized, time-dependent responses of rat cranial bone to repeated mild traumatic brain injuries"

**SUPPLEMENTARY FILES for Dill et al. manuscript entitled *Localized, time-dependent responses of rat cranial bone to repeated mild traumatic brain injuries.***

**SUPPLEMENTARY METHODS**

**ImageJ segmentation protocols**

For mineralized bone reconstruction, to calculate bone volume, surface area, thickness, and diploid volume:

run("8-bit");

run("Statistical Region Merging", "q=10 showaverages 3d");

setAutoThreshold("Otsu dark");

//run("Threshold...");

run("NaN Background", "stack");

run("8-bit");

setAutoThreshold("Otsu dark");

//run("Threshold...");

//setThreshold(8, 255);

//setThreshold(255, 255);

run("Make Binary", "method=Otsu
background=Dark black");

run("Dilate", "stack");

For total tissue (object) volume:

run("8-bit");

run("Statistical Region Merging", "q=10 showaverages 3d");

setAutoThreshold("Otsu dark");

//run("Threshold...");

run("NaN Background", "stack");

run("8-bit");

setAutoThreshold("Otsu dark");

//run("Threshold...");

//setThreshold(8, 255);

//setThreshold(255, 255);

run("Make Binary", "method=Otsu background=Dark black");

run("Dilate", "stack");

run("Remove Outliers...", "radius=40 threshold=30 which=Dark stack");

**SUPPLEMENTARY FIGURE LEGENDS**

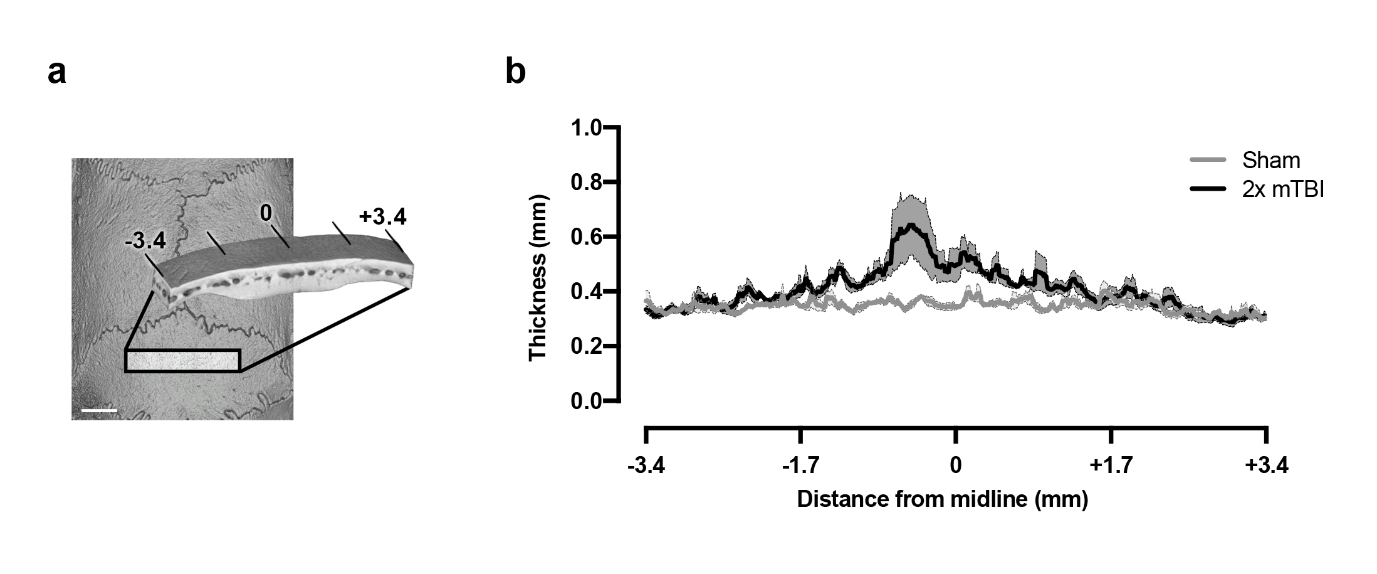

Suppl. Figure 1: The effect of repeated mild TBI on the interparietal bone was greatest near the midline. Quantification of bone thickness across a sagittal cross-section in the interparietal bone at 10 weeks post-injury. Seven hundred microCT images were obtained, spanning -3.4 to +3.4 mm relative to the midline, from five Sham and five 2x mTBI samples. An increase in bone thickness was observed most prominently within approximately 2 mm of the midline. Scale bar = 2 mm. Error bars indicate SEM.

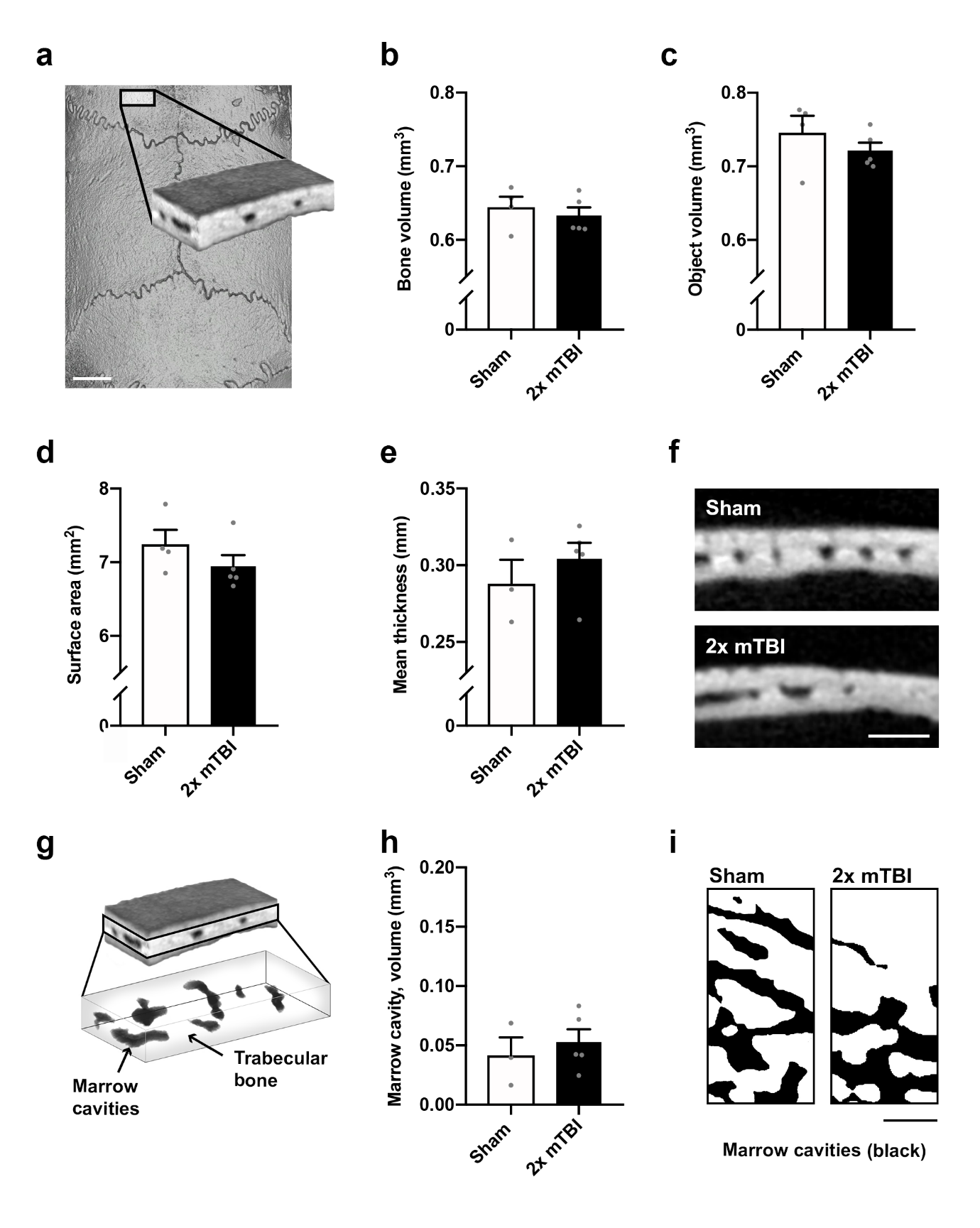

Suppl. Figure 2: Repeated mild TBI did not affect the frontal bone at 10 weeks post-injury. The frontal bone ROI was sampled from the mediocaudal left frontal bone, adjacent to the midline and Bregma suture margins (a; scale bar = 2 mm). There was no significant effect of injury on mineralized bone volume (b; p = 0.54), total object volume (c; p = 0.34), or exterior surface area (d; p = 0.26). Mean mineralized bone thickness, averaged across 100 adjacent sections, was likewise not affected by injury (e; p = 0.40), as demonstrated by representative Sham and 2x mTBI greyscale microCT images from the center of the Z-stack (f; scale bar = 500 µm). Marrow cavities (dark grey) were isolated from trabecular bone (light grey) in the diploë region of the IP bone scan ROIs, for volumetric analysis (g). Total cavity volume was not significantly different between groups (h, p<0.56). Representative superior-view binary projections of the diploë demonstrate comparable marrow cavity (black) density in 2x mTBI mice compared with shams (i; scale bar = 500 µm). N = 5/group. Unpaired student’s t-test.
